# Transcriptomics-based modeling of methionine metabolism effectively estimates sample-wise DNA methylation activity and epigenetic aging

**DOI:** 10.64898/2026.01.30.702880

**Authors:** Tingbo Guo, Pengtao Dang, Yue Fang, Haiqi Zhu, Xiao Wang, Jia Wang, Anjun Ma, Qin Ma, Sha Cao, Chi Zhang

## Abstract

DNA methylation is a central epigenetic modification that regulates gene expression, maintains genomic stability, and guides cellular differentiation. However, direct measurements of DNA methylation, such as whole-genome bisulfite sequencing or DNA methylation arrays, are costly and require substantial DNA input, limiting their scalability for large cohorts and their applicability to emerging modalities such as single-cell and spatially resolved transcriptomics. In this study, motivated by the fact that DNA methylation is fundamentally a metabolic process, we investigate whether sample-wise DNA methylation activity can be inferred directly from transcriptomic profiles of genes involved in methionine and one-carbon metabolism. We show that a compact metabolic model comprising seven core reaction steps and 98 genes accurately predicts total DNA methylation activity across matched transcriptomic and methylation datasets from CCLE, TCGA, GTEx, and an independent single-cell multi-omics data set. Building on this framework, we develop **Total DNA Methylation Activity (TDMA)**, a physics-informed neural network–based score that enables robust estimation of DNA methylation activity from bulk, single-cell, and spatial transcriptomics data. We demonstrate that TDMA captures methylation-dependent transcriptional regulation and identifies genes and pathways under epigenetic control. Applying TDMA to GTEx, we further reveal strong associations between the predicted total methylation activity, chronological aging, and established epigenetic clocks. We also demonstrated that TDMA can serve as a transcriptomics-derived epigenetic clock and highlights age-dependent roles of folate and glutathione metabolism in epigenetic aging. Applying TDMA to single cell and spatial transcriptomics data collected from pancreatic adenocarcinoma (PDAC), we identified that methionine metabolism and DNA methylation regulates T cell cytotoxicity in the tumor microenvironment of PDAC. Together, this work establishes a scalable, modality-agnostic framework for estimating DNA methylation activity from transcriptomics and provides new insights into the metabolic regulation of epigenetic aging.

## INTRODUCTION

DNA methylation is a fundamental epigenetic modification that plays a central role in regulating gene expression, maintaining genomic stability, and guiding cellular differentiation^1^. Aberrant DNA methylation has been implicated in diverse biological processes and diseases, including cancer, aging, and developmental disorders^2^. Consequently, accurate measurement and interpretation of DNA methylation patterns are essential for understanding epigenetic regulation and its contribution to disease mechanisms^3,4^.

Despite substantial technological advances, current experimental approaches for profiling DNA methylations, such as whole-genome or reduced-representation bisulfite sequencing and array-based assays, remain costly and require relatively large amounts of input DNA^5^. These constraints limit scalability to large population cohorts and restrict compatibility with emerging modalities, including single-cell and spatially resolved transcriptomics^6^. As a result, direct methylation measurements are often unavailable precisely in the context where cellular heterogeneity and spatial organization are most biologically informative.

A major use of DNA methylation profiling is to identify genes and pathways under epigenetic control by correlating methylation patterns with gene expression changes^7^. While effective, such analyses depend heavily on direct methylation assays, creating a critical bottleneck for studies that rely solely on transcriptomic data^8^. This gap motivates the development of computational strategies capable of inferring DNA methylation activity from widely available transcriptomic measurements, thereby enabling methylation-aware analyses even when experimental profiling is impractical^9^.

Recent progress in physics-informed neural networks (PINNs) offers a promising framework for estimating metabolic fluxes from steady-state transcriptomic data by embedding biochemical and thermodynamic constraints of metabolic networks into learning algorithms^10,11^. These approaches leverage prior knowledge of reaction stoichiometry and pathway structure to model the nonlinear relationships between gene expression and metabolic activity. Notably, DNA methylation itself is a metabolic reaction within the methionine cycle, in which S-adenosylmethionine (SAM) donates a methyl group to DNA via DNA methyl transferases, producing S-adenosyl homocysteine (SAH)^12^. This biochemical coupling suggests that methylation activity may be inferred from transcriptional states of the surrounding metabolic network.

Motivated by this observation, we hypothesized that the **total DNA methylation activity (TDMA)** of a sample—defined as the aggregate reaction flux of SAM-dependent methylation— can be estimated from transcriptomics using a PINN-based metabolic flux framework. To test this hypothesis, we reconstructed the interconnected methionine, homocysteine, cysteine, glutathione, and folate pathways that govern methyl-group availability and redox balance and developed a message-passing optimization scheme to enable physics-informed flux inference across this network.

Across four independent cohorts with paired transcriptomic and DNA methylation measurements (CCLE, TCGA, GTEx, and a single-cell multi-omics dataset), we demonstrate that sample-wise TDMA can be accurately predicted from transcriptomics alone. We further show that inferred TDMA identifies genes and pathways under methylation-dependent regulation with controlled false discovery rates. Applying the TDMA framework to single-cell and spatial transcriptomics datasets collected from PDAC TME reveal cell-type-specific methylation activity, including associations with T-cell cytotoxicity in the tumor microenvironment and epigenetic heterogeneity across cancer subclones. Moreover, TDMA strongly correlates with chronological aging and established epigenetic clocks, functioning as a transcriptomics-derived surrogate for epigenetic age and uncovering contributions of folate and glutathione metabolism to age-related epigenetic remodeling.

Together, our results establish TDMA as a scalable, assay-independent approach for estimating DNA methylation activity from transcriptomic data and provide a general framework for integrating metabolic modeling with epigenetic analysis. The TDMA predictor is released as an open-source Python package to facilitate broad adoption.

## RESULTS

### Overall consideration and framework of TDMA

The overall design of Transcription-Driven DNA Methylation Activity inference (TDMA) is motivated by two central objectives: (1) determining whether transcriptomic profiles can be leveraged to infer DNA methylation activity, and (2) developing a robust and scalable computational framework for predicting sample-wise DNA methylation levels directly from gene expression data. While methylation at individual CpG sites is influenced by numerous local and context-specific factors, including chromatin accessibility, transcription factor binding, and gene-specific selective pressures, these effects are often difficult to quantify systematically. In contrast, global DNA methylation capacity is primarily constrained by two upstream determinants: the availability of the universal methyl donor S-adenosylmethionine (SAM), produced through methionine metabolism, and the abundance and activity of DNA methyltransferases (DNMTs). These factors collectively govern the overall methylation potential of a cell or tissue.

Based on this principle, TDMA shifts the modeling focus from CpG site–specific methylation to **sample-wise total methylation activity**, which is a more stable and metabolically regulated quantity that can be computationally predicted using transcriptomics data (**Figure 1A**). The central computational hypothesis is that metabolic flux rate through DNA methylation–related reactions can be reliably inferred from the transcriptomic profiles of their associated enzymes. Specifically, estimating flux through the SAM-dependent methylation step provides a quantitative surrogate for global DNA methylation capacity within each sample.

**Figure 1.**
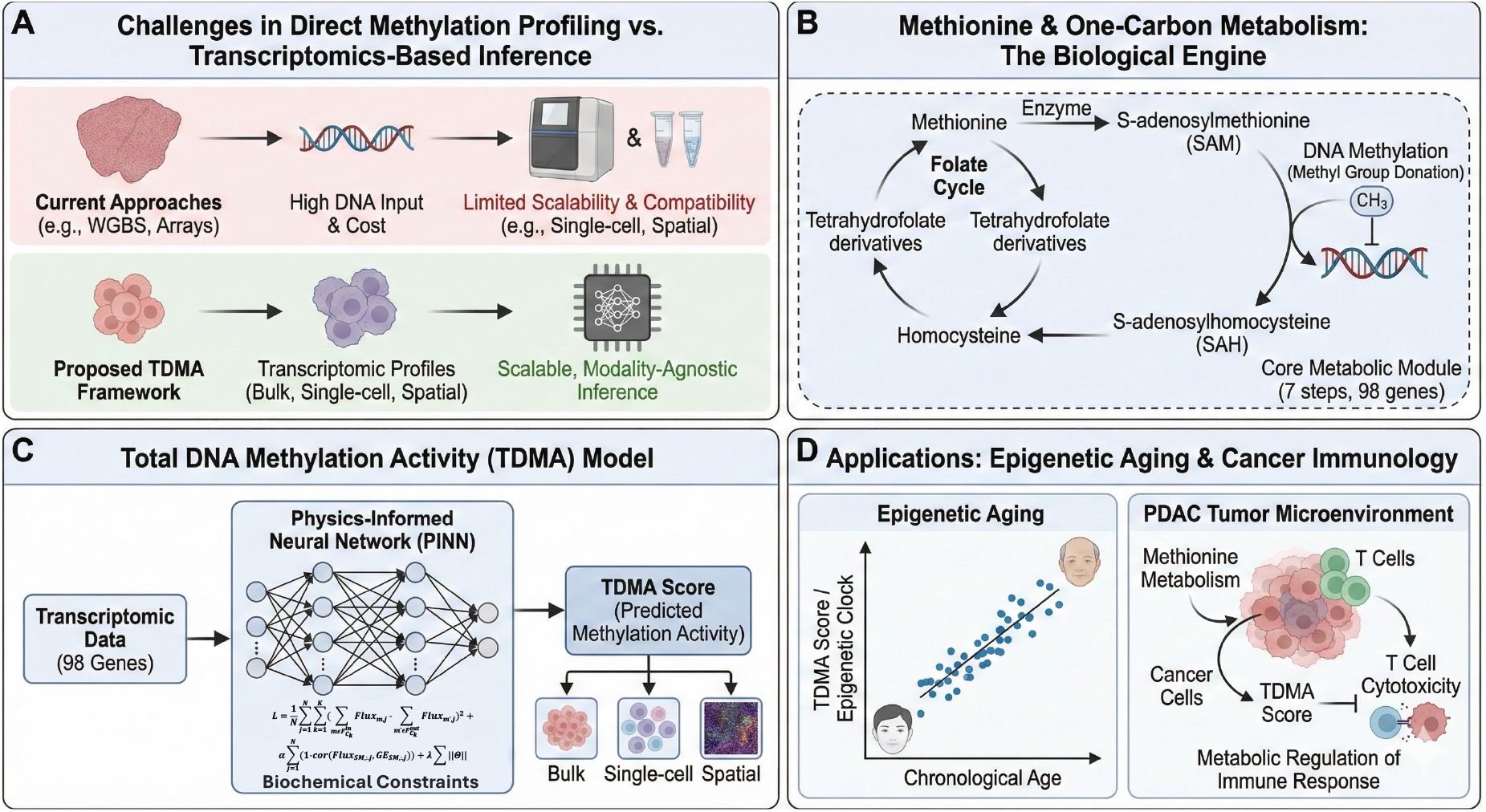
Overview of the TDMA framework. (A) A comparison of experimental and computational assessment of DNA methylation level. (B) Reconstructed methionine and DNA-methylation related modules. (C) PINN model utilized by TMDA. (D) Downstream validations and applications.

To accurately model the DNA methylation activity level, we first reconstructed methionine and DNA-methylation related metabolic network by collecting and curating metabolic reactions and enzymes related to DNA methylation, including methionine, cysteine, homocysteine, and folic acids metabolic pathways and other branches that have significant metabolic exchange with these pathways such as glutathione and purine biosynthesis (**Figure 1B**, see details in **Materials and Methods**). Building upon the networks, we developed a physics-informed neural network framework, termed MPO-SNN (**Figure 1C**), to estimate sample-wise and reaction-level metabolic fluxes from transcriptomic data. MPO-SNN consists of four iterative steps: (1) initializing flux estimates using scFEA, a graph neural network–based flux predictor; (2) projecting predicted fluxes onto the feasible flux-balance solution space to enforce mass conservation; (3) training reaction-specific neural networks that map gene expression of associated enzymes to the projected flux values; and (4) iterating the projection and training steps until convergence. The robustness and accuracy of MPO-SNN were first evaluated using simulated datasets (Supplementary Methods). We first validated the robustness and accuracy of MPO-SNN using simulated data (**Supplementary Methods**). To validate TDMA, we applied the framework to CCLE, TCGA, GTEx, and independent single cell data that have paired transcriptomics and DNA methylation data (**Figure 1D**, see details in **Validation of TDMA using matched DNA methylation and transcriptomics data**). We validated the hypothesis of TDMA and optimized the pathway and flux models that maximize the prediction of DNA methylation in these data sets. In addition, TDMA enables downstream identification of methylation-regulated genes and pathways. Specifically, genes whose expression levels are negatively correlated with predicted methylation activity are prioritized as candidates for methylation-mediated repression, and pathway-level effects are subsequently characterized through enrichment analysis. Together, this framework links cellular metabolism, epigenetic regulation, and transcriptional outcomes within a unified and scalable modeling strategy.

### Mathematical model of TDMA and MPO-SNN algorithm

TDMA approximates metabolic flux rate in Network 1 and 2 using transcriptomics. Given an omics data set *D* and a pathway of *m* reactions, denote the flux rate of the *ith* reaction (*i* = 1, …, *m*) in the *jth* sample in *D* as *F*_*i,j*_. TDMA identifies functions ℱ_*i*_ to estimate *F*_*i,j*_ by *F*_*i,j*_ = ℱ_*j*_ (*D*_, *j*_, Θ^*i*^) and minimizes a loss function 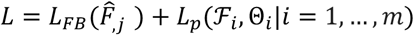, where 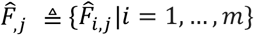 denotes the predicted reaction rates in sample *j*, Θ_*i*_ denotes parameters of ℱ_*i*_, *L*_*FB*_ is a quadratic loss term that regularizes the flux balance of predicted reaction rate, and *L*_*p*_ denotes an aggregated computational loss term to avoid trivial solutions and to conduct variable selection

A key challenge of the above model is to leverage fitting of flux balance condition when training neural networks to approximate the non-linear dependency using gene expression data. To efficiently identify *F*_*i,j*_, we developed a new optimization approach called MPO-SNN (Message Passing-Supervised Neural Network). The MPO-SNN algorithm addresses splitting the fitting process into two iterative steps: (1): Computing 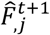 that minimizes 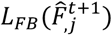 by searching through 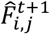 within a certain distance to 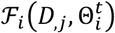 and (2) Supervised Neural Network (SNN) step: Updating 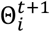 by conducting a supervised training of 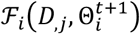 to estimate 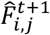, where *t* and *t* + 1 denote successive rounds of iterations. In this approach, the data-fitting step becomes a classic supervised learning problem that can be implemented with additional regularization terms for variable selection or better fitting robustness.

### Validation of TDMA using matched DNA methylation and transcriptomics data

To validate the flux rate of DNA methylation predicted by the TDMA frame can estimate the total DNA methylation level in each transcriptomic sample, we conducted a systematic evaluation using matched DNA methylation and transcriptomics data from three independent sources: the Cancer Cell Line Encyclopedia (CCLE), The Cancer Genome Atlas (TCGA), the Genotype-Tissue Expression (GTEx) project, and a single-cell multi-omics dataset^13–16^. These datasets provide paired gene expressions and DNA methylation measurements across diverse biological systems, including cancer cell lines, primary tumors, normal tissues, and single cells, thereby enabling direct comparison between TDMA-predicted methylation activities and experimentally observed methylation regulation.

We first identified DNA methylation–regulated genes using a well-established epigenetic principle that promoter CpG island methylation is typically associated with transcriptional repression. For each dataset, we computed gene-wise Pearson correlations between measured promoter methylation levels and gene expression across samples and designated genes with strong negative correlations as ground-truth methylation-regulated genes. TDMA is then applied to each data set. To evaluate the agreement between TDMA prediction and experimentally measured DNA methylation level, we computed the Pearson correlation coefficient between (i) TDMA-derived methylation activity and (ii) ground-truth methylation-regulated expression vectors. Higher correlations indicate stronger concordance and greater predictive accuracy.

We first evaluated TDMA using the CCLE dataset, which contains paired gene expression and DNA methylation profiles across a diverse panel of cancer cell lines. Gene expression data were normalized using log_2_(FPKM + 1) and methylation beta values were aligned with matched genes and samples. After quality control, the final dataset comprised 17389 genes across 834 cell lines. Using expression data alone, TDMA identified 8,545 candidate methionine-methylated genes for the representative methylation reaction module. Comparison with experimentally validated methylation-regulated genes derived from paired methylation–expression measurements yielded a Pearson correlation coefficient of **0.7**, indicating strong agreement between predicted and ground-truth gene sets. Scatter plots showed a near-linear relationship along the 45° diagonal (**Figure 2**), further supporting the quantitative consistency of TDMA predictions.

**Figure 2.**
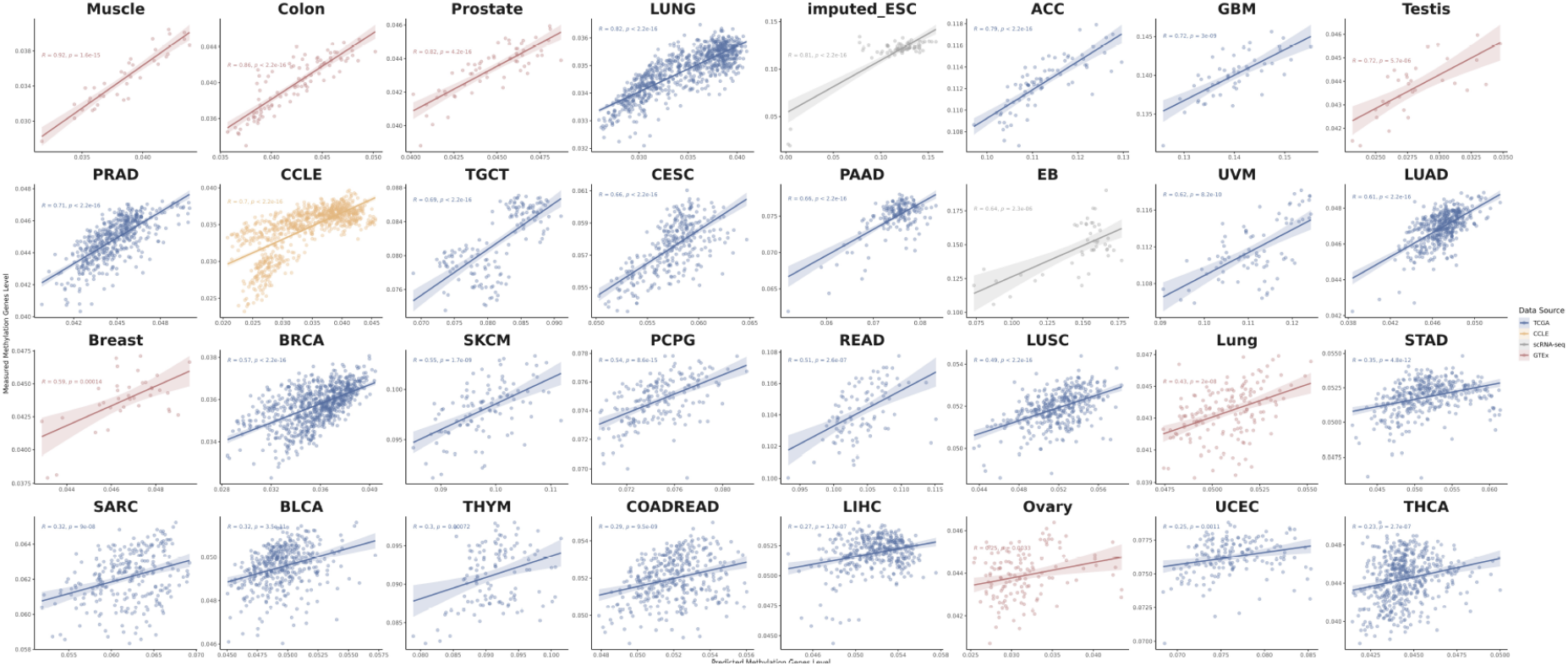
Correlation between TDMA predicted sample-wise total DNA methylation activity and the expression vector of the genes regulated by DNA methylation.

We next assessed TDMA using TCGA multi-omics cohorts spanning eight cancer types (BLCA, COAD, LIHC, UCEC, HNSC, KIRP, KIRC, and THCA). Log_2_(RSEM + 1) normalized expression data was obtained from the Xena platform. Genes were retained if expressed in at least 25% of samples. Across tumor samples, TDMA demonstrated consistently strong performance, with Pearson correlations of 0.82 (LUNG), 0.79 (ACC), 0.72 (GBM), 0.71 (PRAD) and an overall pooled correlation of **0.51**(Figure 2). Normal tissues showed slightly lower but still substantial agreement, with an overall correlation of **0.55**. The modest reduction likely reflects weaker methylation-driven repression and increased regulatory heterogeneity in non-tumor tissues. These results indicate that TDMA accurately predicts methylation regulation across diverse cancer types and disease states.

To further evaluate generalizability beyond cancer systems, we applied TDMA to the GTEx dataset, which provides matched gene expression and DNA methylation profiles across a wide range of normal human tissues. Expression values were log_2_(FPKM + 1) -normalized and processed using the same filtering criteria as TCGA to ensure consistency. TDMA maintained strong concordance between predicted and experimentally validated methylation-regulated genes across tissues. Correlation coefficients remained consistently high across major tissue types (overall correlation = **0.66**; **Figure 2**), demonstrating that TDMA captures fundamental metabolic determinants of DNA methylation that operate broadly in physiological conditions. These results confirm that TDMA is not limited to cancer-specific signals but generalizes to normal tissue homeostasis.

Finally, we evaluated TDMA using a single-cell multi-omics dataset containing matched RNA expression and DNA methylation measurements at single-cell resolution^16^. TDMA-derived methylation activity scores were computed from scRNA-seq profiles and compared against measured methylation levels across individual cells. Despite increased sparsity and technical noise inherent to single-cell data, predicted and observed methylation signals remained strongly concordant (**Figure 2**), demonstrating the robustness of TDMA under high-dimensional and noisy conditions and supporting its applicability at single-cell resolution.

Across four independent platforms, cell lines (CCLE), primary tumors (TCGA), healthy tissues (GTEx), and single cells, TDMA consistently achieved strong concordance between predicted and experimentally validated methylation-regulated genes. These results demonstrate that transcriptomics-driven metabolic inference reliably recapitulates true DNA methylation regulation across biological contexts, establishing TDMA as an accurate, robust, and generalizable framework for inferring methylation activity directly from gene expression data.

### TDMA significantly correlates with epi-genetic age and explains metabolic stresses may facilitate aging-dependent decrease of DNA methylation

DNA methylation has been widely used as a quantitative biomarker of biological aging, forming the basis of numerous epigenetic “clock” models^17–28^. Existing approaches generally fall into two categories: global models that leverage overall DNA methylation levels to estimate epigenetic age, and site-specific models that rely on methylation measurements at selected CpG loci whose effects are largely independent of total methylation burden. Because TDMA accurately infers sample-wise DNA methylation activity directly from transcriptomic profiles, we hypothesized that TDMA-derived methylation estimates could similarly serve as predictors of epigenetic age without requiring direct methylation measurements.

To test this hypothesis, we applied TDMA to the GTEx cohort, a commonly used benchmark for evaluating epigenetic aging models due to its broad tissue coverage and well-annotated age metadata. We computed TDMA-predicted methylation activity for each sample and compared these values with epigenetic ages estimated using twelve established DNA methylation clock algorithms. Across all clocks and tissues, TDMA-inferred methylation activity exhibited consistent negative correlations with predicted epigenetic age (**Figure 3**, M3 column), indicating that lower methylation activity is associated with older epigenetic age. This trend agrees with well-documented observations of global DNA hypomethylation during aging^29,30^.

**Figure 3.**
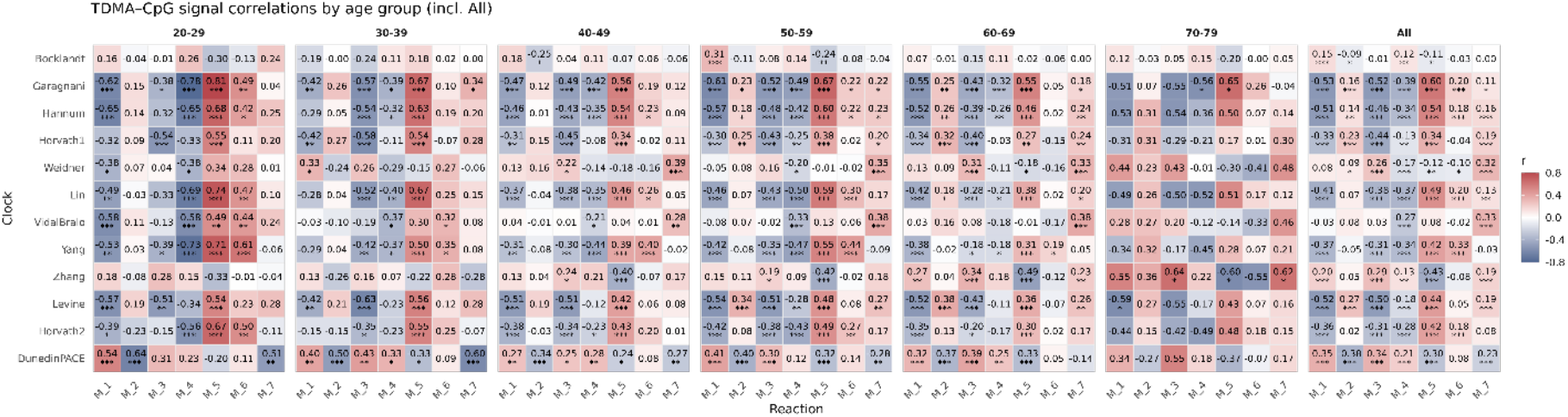
Correlation between epi-genetic age (y-axis) and TDMA predicted DNA methylation activity (x-axis). On x-axis, M1-7 represents the following reactions in Network 2. M1: L-Methionine → S-Adenosyl-L-methionine. M2: S-Adenosyl-L-methionine → L-Methionine. M3: S-Adenosyl-L-methionine → L-Homocysteine. M4: L-Homocysteine + 5,10-Methylenetetrahydrofolate → L-Methionine + Tetrahydrofolate. M5: Folate + Tetrahydrofolate → 5,10-Methylenetetrahydrofolate. M6: 5,10-Methylenetetrahydrofolate → Folate + Thymidine. M7: L-Homocysteine → Glutathione

We next investigated the metabolic mechanisms underlying this age-dependent decline in methylation capacity. SAM, the universal methyl donor for DNA methylation, is synthesized from methionine through one-carbon metabolism, which depends on folate-derived methyl groups. Two major competing branches divert these resources: nucleotide biosynthesis, which consumes folate-mediated one-carbon units, and trans-sulfuration, which converts homocysteine into cysteine and glutathione to support antioxidant defenses. Strikingly, we observed that with increasing age, reduced methylation activity became more strongly associated with elevated antioxidant and glutathione-related fluxes and less associated with folate-mediated methyl-donor metabolism (**Figure 3**, M6–M7 columns).

These findings suggest that age-related oxidative stress may preferentially redirect metabolic resources toward redox homeostasis at the expense of methyl-donor availability, thereby contributing to progressive DNA hypomethylation. Together, these results demonstrate that TDMA not only recapitulates established epigenetic aging signals but also provides a mechanistic metabolic interpretation linking methylation decline to shifts in one-carbon and antioxidant metabolism.

## DISCUSSION

In this study, we present TDMA, a computational framework that enables accurate estimation of sample-wise global DNA methylation activity directly from transcriptomic data. TDMA addresses a longstanding challenge in epigenetics: while DNA methylation is a central regulatory mechanism and a widely used biomarker of development, disease, and aging, direct methylation profiling remains costly, technically demanding, and often unavailable in large or emerging data modalities such as single-cell and spatial transcriptomics. By leveraging metabolic constraints and gene expression signatures of methylation-related pathways, TDMA provides a scalable alternative that infers global methylation capacity without requiring methylation measurements.

Our work makes three key contributions. First, we introduce a physics-informed, message-passing optimization neural network that integrates reconstructed metabolic networks with flux-balance constraints to estimate reaction-level activities underlying DNA methylation. This design combines mechanistic interpretability with the flexibility of neural networks, resulting in improved accuracy and scalability. Second, we demonstrate that TDMA predictions reliably recover methylation-regulated genes and pathways across diverse biological contexts, including cancer cell lines, primary tumors, normal tissues, and single-cell datasets. Third, we show that the framework generalizes across scales, from bulk cohorts to single-cell and spatial transcriptomics, providing a practical and computationally efficient solution for studying epigenetic regulation in large and heterogeneous datasets.

A central conceptual advance of TDMA is the shift from modeling CpG site–specific methylation to estimating **global methylation activity** as a metabolically constrained quantity. While methylation at individual loci is influenced by local chromatin features and stochastic effects, total methylation capacity is primarily determined by systemic metabolic factors, including SAM availability and DNA methyltransferase abundance. By reconstructing methionine, folate, and trans-sulfuration pathways into a unified metabolic map and estimating flux through SAM-dependent methylation reactions, TDMA captures this global regulatory layer. Our analyses demonstrate that flux from methionine to SAM serves as a robust proxy for overall methylation potential, and that transcriptomics-based flux inference can accurately approximate these activities even in the absence of direct methylation measurements.

Beyond methodological advances, our results provide biological insights into the metabolic determinants of epigenetic regulation. For example, TDMA revealed consistent coupling between methylation activity and one-carbon metabolism across cancer and normal tissues, and uncovered age-associated trade-offs between methyl-donor availability and antioxidant pathways. The observed association between reduced methylation activity and increased glutathione synthesis suggests that oxidative stress may redirect metabolic resources away from methylation toward redox homeostasis during aging. These findings highlight the intimate connection between metabolism and epigenetic control and demonstrate how computational flux modeling can reveal mechanistic relationships that are difficult to measure experimentally.

TDMA also enables a broad range of downstream analyses. By providing sample-wise methylation activity scores, the framework supports identification of methylation-regulated genes through correlation analysis, pathway enrichment of epigenetically controlled processes, differential methylation activity across conditions, and spatial mapping of methylation landscapes in tissue sections. These capabilities extend methylation studies to settings where direct methylation profiling is impractical, including very large cohorts, legacy transcriptomic datasets, and high-throughput single-cell or spatial technologies.

Several limitations should be noted. First, TDMA estimates global methylation capacity rather than locus-specific methylation states and therefore cannot replace targeted CpG-level assays when precise site resolution is required. Second, while TDMA focuses on DNA methylation as a primary epigenetic outcome, methyl-group metabolism regulates a broader spectrum of methylation-dependent processes beyond DNA alone. S-adenosylmethionine (SAM) also serves as the universal methyl donor for RNA, histone, and protein (amino-acid) methylation, which collectively influence chromatin organization, transcriptional regulation, and post-transcriptional control. Extending the current framework to jointly model these additional methylation substrates may provide a more comprehensive view of cellular methylation demand and improve estimation of global methylation dynamics. Future work will therefore expand TDMA to incorporate multi-layer methylation processes, enabling unified modeling of DNA, RNA, and histone methylation within a systems-level metabolic–epigenetic framework.

In summary, TDMA provides a mechanistically interpretable and computationally scalable framework for inferring DNA methylation activity directly from transcriptomic data. By bridging metabolism and epigenetic regulation, our approach expands the accessibility of methylation analysis to diverse data types and large-scale studies, opening new opportunities to investigate how metabolic states shape gene regulation in development, aging, and disease.

## MATERIALS AND METHODS

### Reconstruction of DNA methylation-related metabolic pathway

To enable reliable and mechanistically grounded prediction of DNA methylation, we systematically curated and reconstructed a DNA methylation–centered metabolic network through extensive integration of reactions and annotations from KEGG, Recon3D, Reactome, and prior literature reports^31–34^. Reactions were manually verified and harmonized across databases to ensure biochemical consistency and gene–enzyme mapping. This effort yielded a compact yet comprehensive metabolic module capturing the biochemical processes that regulate methyl-donor availability, nucleotide biosynthesis, and cellular redox homeostasis.

Based on this annotation, we constructed **Network 1** that comprises 24 reactions with associated enzymes and metabolites. The network explicitly models the coordinated interplay among methionine metabolism, one-carbon/folate metabolism, trans-sulfuration, and nucleotide synthesis pathways, which collectively determine cellular DNA methylation potential. These pathways compete for and recycle shared one-carbon and sulfur resources, thereby coupling epigenetic regulation to metabolic state.

The network initiates with methionine adenosyltransferases (MAT1A/MAT2A/MAT2B), generating S-adenosylmethionine (SAM), the universal methyl donor for DNA methylation. SAM-dependent methyl transfer reactions, catalyzed by DNA methyltransferases (DNMT1, DNMT3A, DNMT3B, DNMT3L), convert SAM to S-adenosylhomocysteine (SAH). SAH is subsequently hydrolyzed by AHCY, AHCYL1, and AHCYL2 to regenerate homocysteine, completing the methionine cycle. Sulfur derived from methionine is further redistributed through enzymes such as LIAS, linking methylation metabolism to downstream sulfur utilization.

To account for alternative sinks and recycling routes that influence methyl-donor availability, the network incorporates polyamine biosynthesis (AMD1, SRM) and the methionine salvage pathway (MTAP), which recycle methylthioadenosine (MTA) back to methionine. Additional enzymes, including IL4I1 and TAT, were included to represent side pathways that modulate methionine flux and therefore methylation capacity. The one-carbon metabolism module integrates the folate cycle with methionine recycling. Remethylation of homocysteine to methionine is mediated by BHMT and MTR using folate-derived methyl groups. SHMT1/2 and MTHFR generate and redistribute one-carbon units from serine, supplying substrates required for both nucleotide synthesis and methylation reactions. These shared intermediates create a direct metabolic trade-off between DNA methylation and anabolic demands. Finally, the trans-sulfuration pathway (CBS, CTH) channels homocysteine toward cysteine and glutathione (GSH) biosynthesis. Downstream antioxidant enzymes (GCLC, GSS, GPX1–8, GSR) regulate redox balance, thereby linking methylation capacity to oxidative stress. Reactions supporting nucleotide synthesis (e.g., DHFR, MTHFD1, MTHFD2) and purine metabolism (NT5C, NT5E) were also incorporated to model competition for one-carbon units, further constraining methyl-group availability for DNA methylation.

Collectively, Network 1 provides a mechanistic representation of the metabolic determinants of DNA methylation, explicitly coupling methyl-donor generation, recycling, and competing biosynthetic demands. This reconstruction forms the biochemical foundation for subsequent quantitative modeling and prediction of methylation-related metabolic states.

To facilitate quantitative modeling and improve computational tractability, we further derived a reduced representation of Network 1 by collapsing sequential reactions with tightly coupled fluxes into higher-level composite reactions. This reduction strategy preserves overall mass balance and biochemical functionality while decreasing network dimensionality and minimizing redundancy introduced by intermediate metabolites. Reactions operating in linear or near-linear cascades were therefore lumped into single effective steps, resulting in a compact coarse-grained network, referred to as **Network 2**.

Network 2 summarizes the methylation-centered metabolism into seven functional modules that capture the dominant flow of carbon and sulfur through methionine activation, methyl transfer, one-carbon recycling, folate interconversion, nucleotide synthesis, and antioxidant production. The resulting composite reactions are:

1. Methionine activation: L-Methionine → S-Adenosyl-L-methionine
2. SAM turnover / methyl utilization: S-Adenosyl-L-methionine → L-Methionine
3. SAM demethylation to homocysteine (methylation flux): S-Adenosyl-L-methionine → L-Homocysteine
4. Homocysteine methylation: L-Homocysteine + 5,10-Methylenetetrahydrofolate → L-Methionine + Tetrahydrofolate
5. Folate interconversion: Folate + Tetrahydrofolate → 5,10-Methylenetetrahydrofolate
6. Thymidine/nucleotide synthesis branch: 5,10-Methylenetetrahydrofolate → Folate + Thymidine
7. Trans-sulfuration and antioxidant production: L-Homocysteine → Glutathione

By explicitly aggregating these reactions into functional modules, Network 2 captures the principal metabolic trade-offs governing DNA methylation capacity: (i) SAM generation and utilization, (ii) recycling of homocysteine through the folate-dependent one-carbon cycle, and (iii) competition between methylation, nucleotide biosynthesis, and glutathione synthesis for shared metabolic resources. Compared with Network 1, this reduced network substantially lowers model complexity while retaining the essential biochemical constraints required for downstream flux estimation and predictive analysis.

